# Microbial ecology of *Ixodes scapularis* from Central Pennsylvania, USA

**DOI:** 10.1101/2020.07.11.197335

**Authors:** Joyce M. Sakamoto, Gabriel Enrique Silva Diaz, Elizabeth Anne Wagner

## Abstract

(1) Background: Native microbiota represent a potential resource for biocontrol of arthropod vectors. *Ixodes scapularis* are mostly inhabited by the endosymbiotic *Rickettsia buchneri,* but the bacterial communty composition varies with life stage, fed status, and/or geographic location. We investigated sex-specific bacterial community diversity from *I. scapularis* collected from central Pennsylvania between populations within a small geographic range. (2) Methods: We sequenced the bacterial 16S rRNA genes from individuals and pooled samples and investigated the abundance or infection frequency of key taxa using taxon-specific PCR and/or qPCR. (3) Results: Bacterial communities were more diverse in pools of males than females. When *R. buchneri* was not the dominant taxon, Coxiellaceae was dominant. We determined that the infection frequency of *Borrelia burgdorferi* ranged between 20 to 75%. Titers of *Anaplasma phagocytophilum* were significantly different between sexes. High *Rickettsiella* titer in pools were likely due to a few heavily infected males. (4) Conclusion: Bacterial 16S sequencing is useful for establishing the baseline community diversity and focusing hypotheses for targeted experiments. However, care should be taken not to overinterpret data concerning microbial dominance between geographic locations based on a few individuals as this may not accurately represent the bacterial community within tick populations.

## Introduction

Ticks are obligately hematophagous arachnids that are found worldwide parasitizing vertebrates. Only eight of the 84 described tick species in the contiguous United are responsible for the majority of tick-borne disease (TBD) [1]. Reported cases of TBD have more than doubled since 2004 and account for 76.5% of total vector-borne disease cases [2].

In the United States, the vector of the Lyme disease pathogen *Borrelia burgdorferi* is the blacklegged tick *Ixodes scapularis. I. scapularis* also transmits other parasites that cause babesiosis, anaplasmosis, ehrlichiosis, and Powassan encephalitis [3]. Methods of disease prevention (landscape management, acaricides, repellents, tick checks) can be effective, but are not 100% efficient and are often difficult to do when pets regularly visit the outdoors or humans work in high tick-load areas. Vaccines against some tickborne diseases offer protection, but there is no currently available Lyme vaccine for humans, although there are some promising candidates on the horizon [4,5].

Alternative approaches to arthropod vector control may include the manipulation of microbial communities to identify possible antagonists or targets for tick and/or pathogen control. Microbial communities may modulate the invasion, replication, and/or transmission of pathogens in vector arthropods or potentially inhibit transmission to vertebrate hosts[6–9]. Endosymbiotic bacteria can also influence the biology of the arthropods themselves, altering life history characteristics positively or negatively [10– 12]. Several comprehensive reviews of the effects of symbiotic bacteria of ticks have shown that, while some bacteria have co-evolved with ticks, others may have been acquired independently [13,14]. The exact role of these taxa are not entirely understood—while many are transgenerationally transmitted, others may be acquired through the soil or leaf litter in which the ticks spend a great majority of their lives [15– 17]

Native microbiota (including obligate symbiotic bacteria) are attractive targets for nonchemical vector control strategies. One paratransgenic strategy (Cruzigard) utilized transgenic actinomycete *Rhodococcus rhodnii* to target the Chagas disease parasite *Trypanosoma cruzi,* but was not implemented in full field trials, perhaps in part due to anti-GMO sentiment[18,19]. In contrast, the World Mosquito Program (which releases *Wolbachia-infected* mosquitoes to control Dengue virus) has been accepted and implemented in 12 different countries [20]. Care and thorough investigation must be taken in choosing candidate bacteria before implementation. The introduction of nonnative bacteria can reduce pathogen transmission in one vector species but may increase transmission in another, may adversely affect/be affected by the native microbiota, or alter the vector biology to indirectly affect vectorial capacity [8,21,22].

Multiple approaches have been used to identify the key players in the microbial communities of ticks. In earlier studies, 16S sequencing of agar-grown bacterial colonies identified culturable taxa, and PCR primers/antibodies specific to known taxa were useful for determining infection frequencies [23–26]. Next-generation sequencing platforms offered deeper coverage of the microbial community, promising both to identify unculturable organisms and to detect rare taxa [27,28]. Sequencing fragments of the eubacterial 16S rRNA gene represents a powerful way to assess variation at the individual and population level (through sample pooling). Testing pools containing multiple individuals allows one to screen more populations at a cost of losing resolution at the individual level. Ideally, one needs to strike a balance between sample analysis cost and broadness of sampling to gain an accurate picture of microbiome variation across populations.

In this study, we used 16S rRNA sequencing to 1) determine the baseline microbial diversity in tick populations within a relatively small geographic area, 2) confirm the species identity of key taxa using taxon-specific PCR and Sanger sequencing, and 3) estimate the relative abundance of key bacterial taxa by PCR and/or qPCR in pooled DNA and individual ticks collected from central Pennsylvania.

## Methods

### Tick collections

Adult male and female *I. scapularis* were collected from central Pennsylvania from 2012 to 2019. The collection sites were separated by as little as 0.804 to as much as 43.13 km (0.5-26 miles) apart (Figure 1). Field collections were conducted using a drag cloth (36” x 45”; 91.44 cm x 114.3 cm). Samples were stored alive in 20 ml scintillation vials until returned to the laboratory for immediate surface sterilization by washing in 70% ethanol for 15 seconds, then 1 minute in 10% bleach, followed by three sequential washes in autoclaved, nuclease-free water, dried on autoclave-sterilized filter paper, and stored at −80 °C until processed for DNA extraction. Samples were sorted to sex and species confirmed before extraction [29].

**Figure 1.**
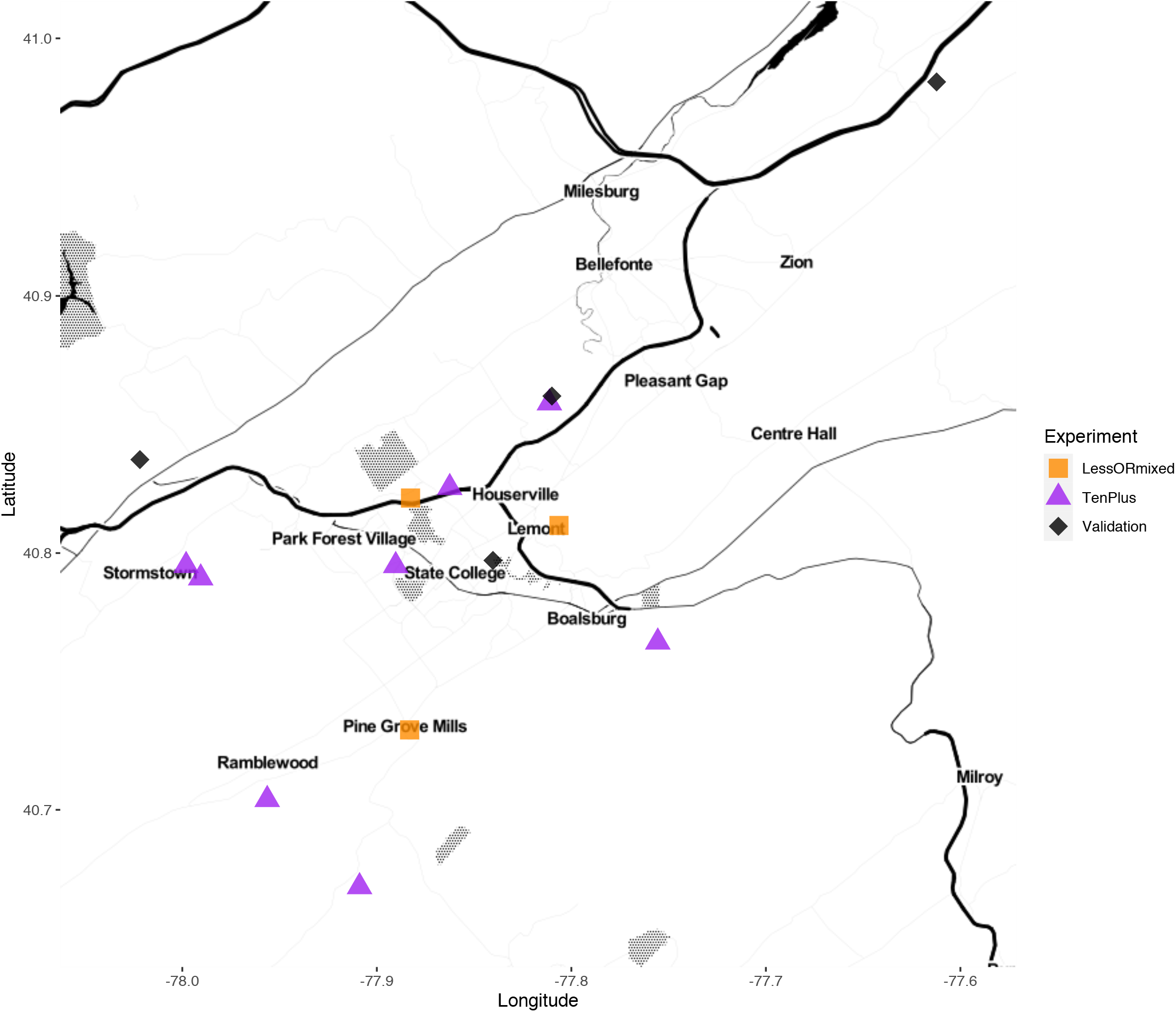
Field collection sites. Squares (“LessORMixed”) represent pools containing less than 10 individual ticks, mixed pools of male and female, or both. Triangles (“TenPlus”) represent pools containing 10 or more individuals. Diamonds (“Validation”) represent collection sites assayed by qPCR.

### 16S rRNA sequencing of individually extracted ticks

For preliminary 16S rRNA sequencing, genomic DNA was extracted from thirty individual adult ticks collected in 2012 (Table 1). DNA was extracted from each tick using the DNeasy Blood and Tissue kit (#69506, Qiagen, Germantown, MD). Samples were submitted for 16S rRNA sequencing of the hypervariable region v6 on the Illumina MiSeq platform to assess the baseline species composition of field-collected blacklegged ticks.

**Table 1.** Collection information for tick samples sequenced by Illumina MiSeq. All samples were collected from wild populations (from central Pennsylvania and 2 pools from Mississippi). Samples were either individually sequenced or sequenced as pools of DNA extracted from individuals. Hypervariable regions sequenced are included.

| Collection year | SampleID | Population | Sex | Individuals or Pooled (N) | V4 or V6 |
| --- | --- | --- | --- | --- | --- |
| 2012 | BHF01-F10 | BigHollow (2013seq) | Female | Individuals (10) | V6 |
| 2012 | BHM01-M10 | BigHollow (2013seq) | Males | Individuals (10) | V6 |
| 2012 | SCF01-F10 | Shaver's Creek (2013seq) | Females | Individuals (10) | V6 |
| 2013 | MB001 | Stormstown | Female | Pooled (20) | V4/V6 |
| 2013 | MB002 | Stormstown | Males | Pooled (20) | V4/V6 |
| 2013 | MB004 | BlueCourse | Female | Pooled (18) | V4/V6 |
| 2013 | MB003 | BlueCourse | Males | Pooled (15) | V4/V6 |
| 2013 | MB006 | APDS | Female | Pooled (16) | V4/V6 |
| 2013 | MB005 | APDS | Males | Pooled (14) | V4/V6 |
| 2013 | MB008 | Big Hollow | Female | Pooled (20) | V4/V6 |
| 2013 | MB007 | Big Hollow | Males | Pooled (15) | V4/V6 |
| 2013 | MB009 | Galbraith | Female | Pooled (20) | V4/V6 |
| 2013 | MB010 | Galbraith | Males | Pooled (20) | V4/V6 |
| 2013 | MB012 | Benner Springs | Female | Pooled (20) | V4/V6 |
| 2013 | MB011 | Benner Springs | Males | Pooled (20) | V4/V6 |
| 2013 | MB013 | SGL176 | Female | Pooled (20) | V4/V6 |
| 2013 | MB014 | SGL176 | Males | Pooled (20) | V4/V6 |
| 2014 | MB015 | Shaver's Creek | Female | Pooled (20) | V4/V6 |
| 2014 | MB016 | Shaver's Creek | Males | Pooled (20) | V4/V6 |
| 2014 | Lem | Lemont | Female | Pooled (3F) | V4/V6 |
| 2014 | PGM | Pine Grove Mills | Mixed | Pooled (4F, 4M) | V4/V6 |
| 2014 | Toft | Toftrees | Mixed | Pooled(11F, 9M) | V4/V6 |
| 2009 | MS1 | Mississippi clade 1 (F3, F5, G4, G6)<sup>ce</sup> | Females | Pooled (4) | V4/V6 |
| 2009 | MS2 | Mississippi clade 2 (F6, F8, G11)<sup>de</sup> | Females | Pooled (3) | V4/V6 |

<sup>a</sup> Individual males from this population/sampling time did not have high enough DNA quality for sequencing
<sup>b</sup> Mississippi pool#1 included the following females: F3 (MS\_1.F2, Copiah Co, MS, 17 Nov 2006, JGoddard); F5(MS\_3.F1, Copiah Co, MS, 28 Dec 2006, JGoddard); G4 (MS\_7.F3, Copiah Co, MS, 29 February 2008, JGoddard); and G6 (MS\_8.F1, Copiah Co, MS, 19 November 2008, JGoddard).
<sup>c</sup> Mississippi pool#2 included the following females: F6 (MS\_4.F1, Copiah Co, MS, 10 January 2007, JGoddard), F8 (MS\_4.F3, Copiah Co, MS, 10 January 2007, JGoddard), and G11 (MS\_6.F1, Copiah Co, 4 April 2007, JGoddard)

### 16S rRNA Sequencing of pooled DNA from individually extracted ticks

After the preliminary sequencing run on individual tick DNA, we chose to expand to more populations, opting to pool DNA from individually extracted ticks due to budget constraints. This approach allowed us to use the sequence results from the pools to then target specific taxa and determine infection frequency or titer, depending on the question.

A total of 298 ticks (collected in 2013-2014) were split by sex across 8 sites (=16 pools) and used for Illumina sequencing and qPCR validation (Table 1). Each sample was surface-sterilized as described above, allowed to air-dry on autoclave-sterilized filter paper in a clean plastic petri dish, and bisected longitudinally with a flame-sterilized new razor blade. One half was used for extraction using the DNeasy Blood and Tissue kit (Qiagen) or the GenElute Bacterial DNA extraction kit (#NA2110, Millipore Sigma, St. Louis, MO). The other half was archived at −80 °C.

An additional 38 ticks representing 5 pools were also tested. These samples were pooled before DNA extraction (i.e. not individually extracted) from populations and represented either mixed samples of males and females or populations with less than 10 individuals. These ticks were surface-sterilized and extracted as described above, but were not bisected individually, nor were they archived. For comparison to geographically distant populations, we included two pools of extracted DNA from individual female ticks representing two mitochondrial clades from Mississippi described previously [30], (Table 1).

Genomic DNA for each tick was quantified with a Nanodrop spectrophotometer and adjusted to at least 5 ng/ul. Equal volumes of adjusted DNA were pooled by population and sex and submitted for sequencing (for a total of 16 submitted pools). The extracted genomic DNA was submitted for Illumina MiSeq sequencing using primers to amplify either the V6 or V4 hypervariable region of the bacterial 16S rRNA gene region.

### Analysis of 16S rRNA sequences

Sequences were aligned, filtered to remove chimeric sequences, and analyzed using Dada2 and visualized using the R packages Phyloseq, Microbiome, and ggpubr [31– 34]. Reads were assigned taxonomic identity using the Dada2 taxonomy assigner and either the Greengenes or the Silva (v128) reference database of eubacterial 16S rRNA [35,36]. Unique amplicon variants were filtered to exclude zeros and singletons, so the Chao index was not appropriate for the data. Shannon and inverse Simpson indices were used for measuring richness and evenness. Statistical comparisons between groups were performed using the Kruskal Wallis test. Beta diversity was quantified the Bray-Curtis matrix. Principal coordinate analyses (PCoA) were calculated and plotted to visualize bacterial community structure between groups using Phyloseq. Statistical comparison between groups was performed to run permutational multivariate analyses of variance (PERMANOVA). The significance level was set to 0.05.

### Taxon-specific amplification and sequence confirmation

We used taxon-specific primers (either previously published or designed for this study) on a subset of tick samples positive for *Rickettsia, Rickettsiella, Borrelia,* and *Cardinium* (Tables 2, 3). Amplicons were separated on a 0.5x TAE-2% agarose gel. Bands were excised and purified for cloning with the StrataClone PCR Cloning kit (#240205, Agilent, Santa Clara, CA), following manufacturer’s guidelines. Plasmids were purified using the E.Z.N.A. Plasmid mini kit (Omega Biotek #D6942) and sequenced in both directions on an ABI 3130/Genetic Analyzer. Sequences were compared to known sequences in the Genbank NR nucleotide database.

Sequences were amplified with species- or genus-specific primers (Table 2) and were aligned, trimmed, and phylogenetically analyzed in MEGA7 [48]. The evolutionary history was inferred by using the Maximum Likelihood method based on the General Time Reversible model[49]. Bootstrap consensus was inferred from 1000 replicates and branches in fewer than 50% bootstrap replicates were collapsed [50]. Genbank Accession numbers used for each taxon/gene analysis are listed in Table 4.

**Table 2.**
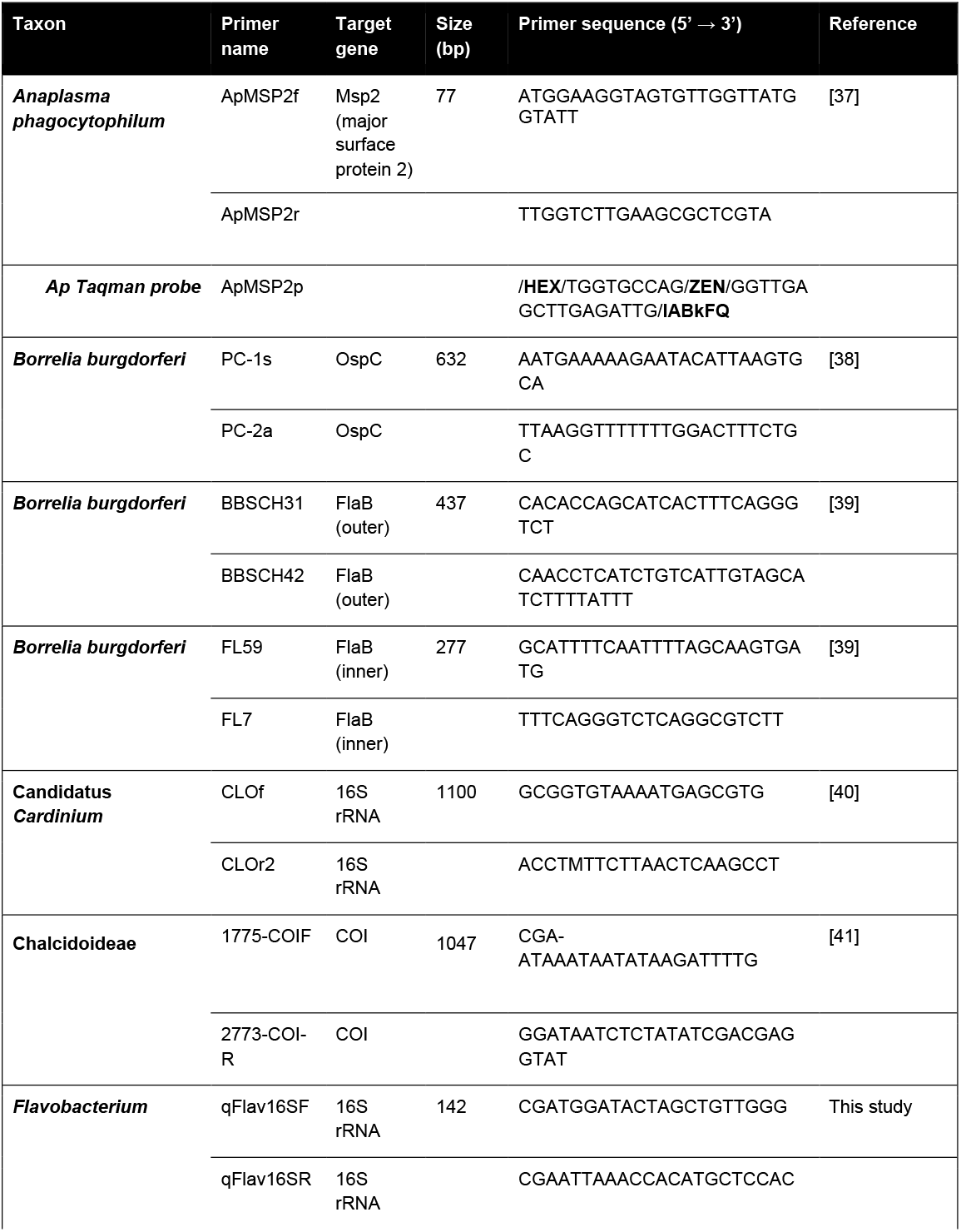
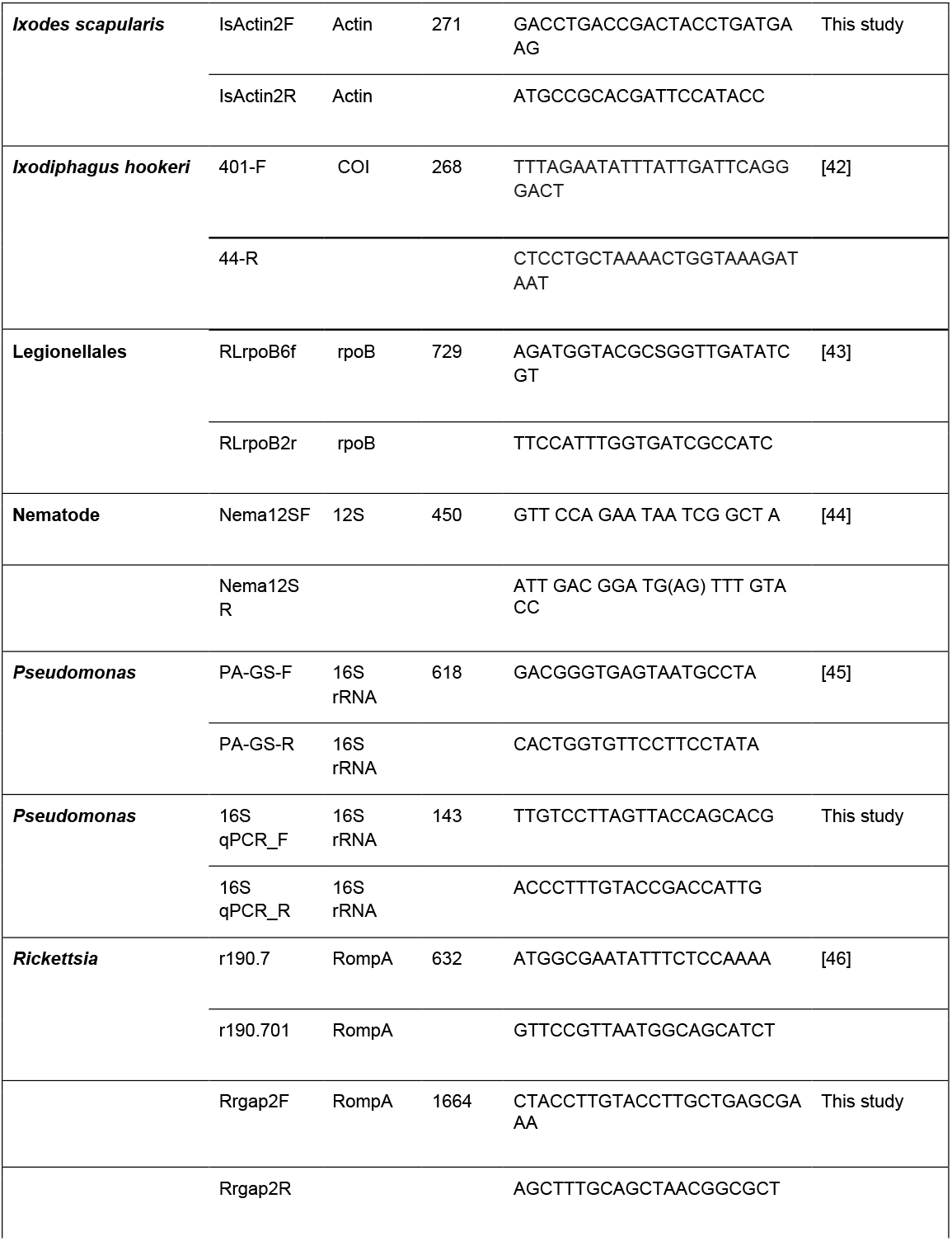
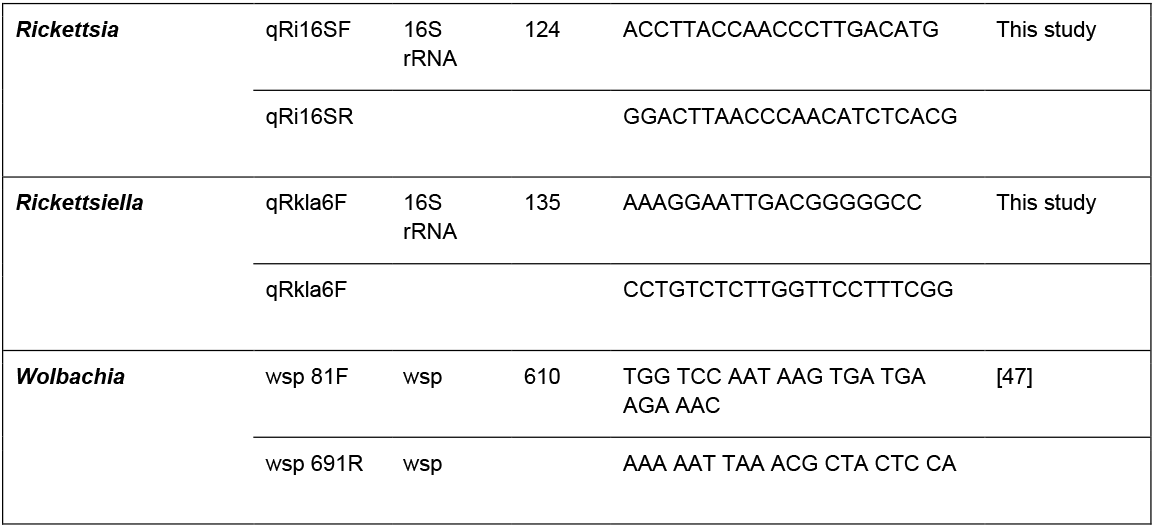
Oligonucleotides used in this study.

**Table 3.**
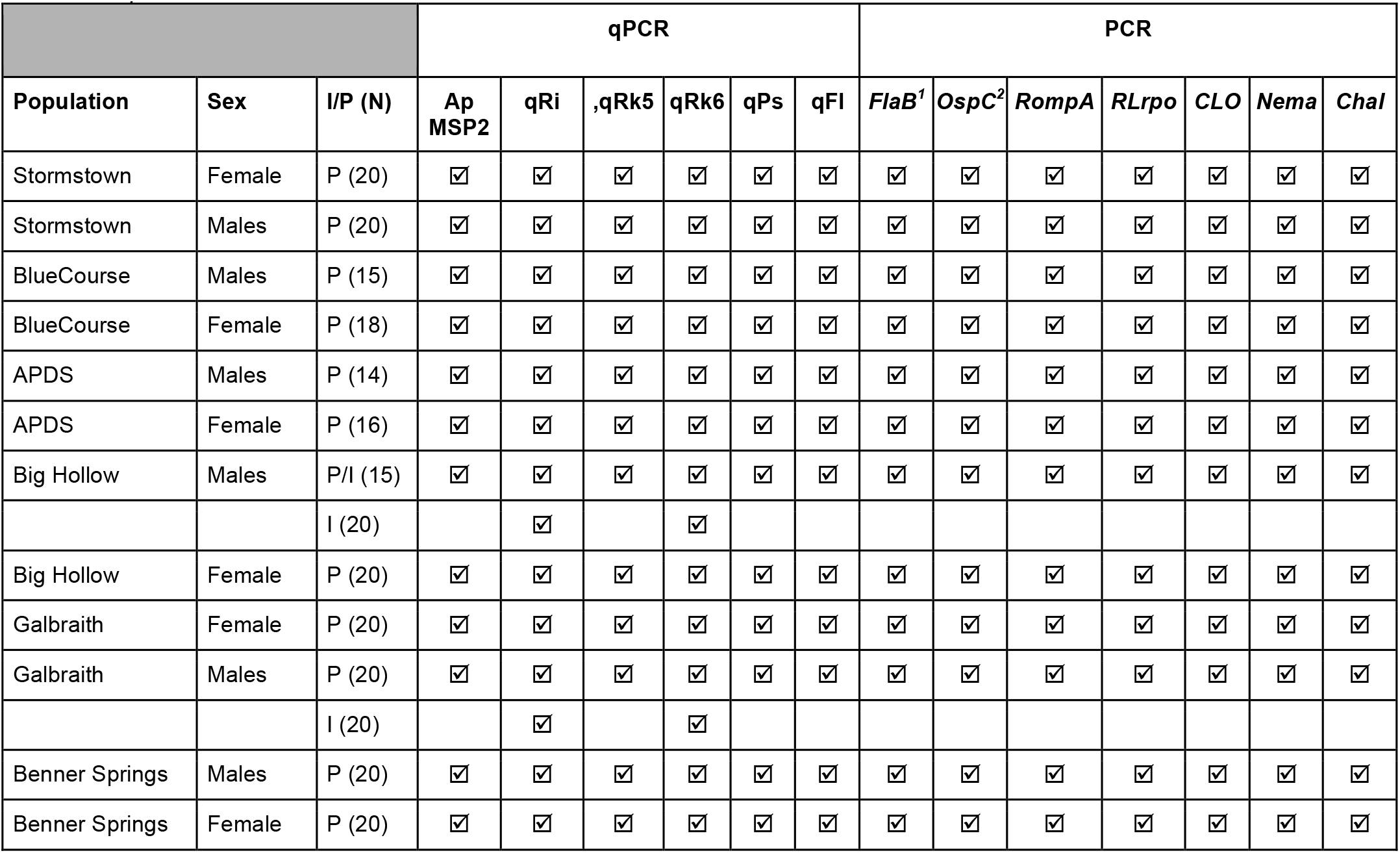
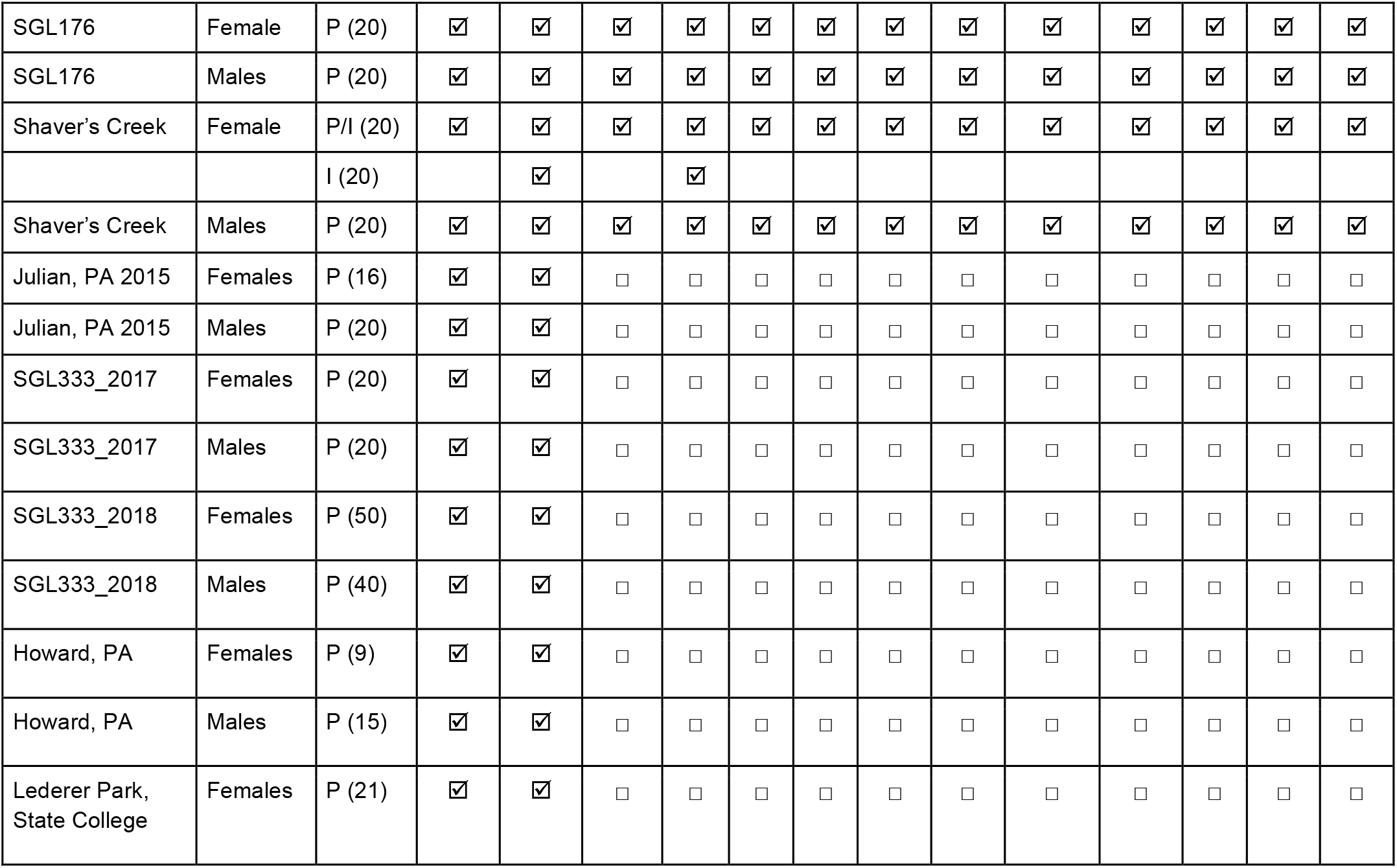
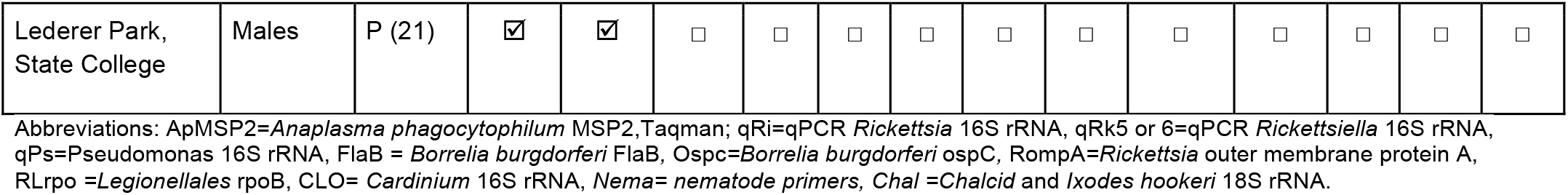
Samples used for validation. All samples were collected from wild populations (from central Pennsylvania). Samples were assayed as individuals (I) or pooled DNA (P) by PCR or qPCR. Housekeeping gene used for qPCR was *Ixodes scapularis* actin2.

**Table 4.** Genbank Accession Numbers used in Phylogenetic Analysis

| **Taxon** | **Gene** | **Genbank Accession Number** |
| --- | --- | --- |
| ***Rickettsia*** | **RompA** | JFKF01000169.1, LN794217.1, CP003340.1, CP047359.1, CP003375.1, CP040325.1, CP001612.1, CP003308.1, CP001227.1, CP012420.1, CP000683.1, CP003342.1 |
| ***Rickettsiella*** | **rpoB** | KP985349.1, KP985348.1, KP985347.1, KP985346.1, KP985354.1, KP985353.1, EF694042.1, JF288929.1, KP985351.1, KP985350.1, AP018005.1, KP985357.1, KP985356.1, KP985355.1 |
| ***Cardinium*** | **16S rRNA** | AB241135.1, AB241129.1, GU731426.1, AB241131.1, AB241130.1, AY753169.1, MH057615.1, AY635291.1, AB001518.1, LN829689.2, JN204482.1, LC159289.1, AB241132.1, AF350221.1, AB116514.1, KX022134.1, AB506775.1, AB506773.1, AB506774.1, GQ206320.1, CP022339.1 |

Although *Wolbachia* has been described from *I. scapularis,* its presence could be as an obligate symbiont of the ticks, or it could be due to the presence of tick endoparasites such as parasitoid wasps (e.g. *Ixodiphagus hookeri)* or nematodes [42,51]. Therefore, we assayed the pooled DNA with PCR specific to *Wolbachia,* nematode or chalcid wasp genes (Tables 2–3).

### *Borrelia burgdorferi* infection frequency

Sex-specific infection frequencies from each population were assessed by amplifying the *Borrelia burgdorferi* OspC gene (non-nested PCR) and the flagellin FlaB gene (nested PCR) following conditions described previously [38,39](Table 2, Table 3). DNA extracted from individual ticks was tested from each of the pooled populations and the percentage of infected individuals per total pool (sex-location) was calculated.

### Taxon-specific quantitative PCR

Taxon-specific qPCR primers were used to assess relative titer (Table 3) on individuals or pools with at least 10 individuals or more per sex-population pool. In addition to running assays on pools from which we had Illumina sequence data, we tested the *Rickettsia* and *A. phagocytophilum* qPCR assays on pooled DNA from 232 ticks (10 pools of males or females) collected from other locations and/or from other years (Table 3).

qPCR primers were designed for *Rickettsia, Flavobacterium, Pseudomonas,* and *Rickettsiella* using the NCBI tool Primer-blast [52] and confirmed to match the desired taxa by Sanger sequencing of the amplicons. PCR mixes were made with the PerfeCTa SYBR Green FastMix (Quanta Bioscience, Inc., Gaithersburg, MD). All reactions were run on a Rotor-Gene Q 5plex HRM System (QIAGEN) for 40 cycles x (95 °C/10s, 60 °C/15s, 72 °C/15s). Relative titer for each of gene was compared to the reference gene *I. scapularis* actin gene (Genbank accession # XM_029983741.1). Assays run before v4 sequencing (on *Rickettsia, Flavobacterium,* and *Pseudomonas* against Actin) were run in duplicate. After v4 sequencing, all qPCRs were run in triplicate *(Rickettsia, Rickettsiella,* and actin).

We used a Taqman assay for detection of *Anaplasma phagocytophilum* [37], with the following modifications: we used a PrimeTime® Standard qPCR Assay (in which primers and probes are mixed and received in a single tube) (Integrated DNA Technologies, Inc., Iowa, USA) and the probe with a double-quenching Zen/Iowa Black instead of TAMRA. Relative titer for each of gene was compared to the reference gene *I. scapularis* actin gene. Reactions were run for 40 cycles x (95 °C/15s, 60 °C/60s). Each sample was run in triplicate.

## Results

### Individual vs. pool (hypervariable region v6)

The relative abundance of bacterial taxa from individual tick samples suggested that the microbial community composition of females from Shaver’s Creek was more similar to males from Big Hollow (BHM) (Supplementary Figure 1). We observed that the most abundant taxa were Rickettsiaceae, and Pseudomonadaceae (Supplementary Figure 1). We chose to evaluate the titers of *Rickettsia, Pseudomonas,* and *Flavobacterium* by qPCR. We found the titers of the latter two taxa to be considerably less than indicated by sequence counts, although we did detect a significant sex-specific difference in these three taxa (Supplementary Figure 2). We did not test the effects of sex and population as these could not be accurately evaluated with the limited number of samples and with only one pool of males.

The individual ticks represented a preliminary sequencing run. We originally chose to sequence the v6 region because at the time we thought it was better suited for genus and species-level resolution. We had Illumina data on 30 individuals from two populations but found that the amount of variability from one pool of females (Shaver’s Creek) was very different from the females from another pool (Big Hollow) and more closely matched that of the males. Further, we did not detect Coxiellaceae, but found that when Rickettsiaceae was in low abundance, a Pseudomonadaceae was more abundant. These data represented only two populations, so we revisited these populations to collect fresh ticks for the pooled sequencing run in addition to 6 other populations. We found that this time, Pseudomonadaceae dominated not only every one of our pools but also that of every sample (from unrelated experiments) in the same sequencing run. We suspected that there was a v6 primer bias and re-sequenced the v4 region of the same pools for subsequent analyses. We also suspected that Greengenes was not an appropriate reference database for environmental data and ran subsequent taxonomic assignments with the Silva database.

### Taxonomic assignment and taxa summary of pools (v4)

We compared the 21 different pools (16 paired male and female pools plus 5 pools of mixed-sex or fewer than 10 samples/pool) and observed that the diversity between groups was similar except for one female pool from population “APD”. We used Grubb’s outlier test to determine that this was an outlier pool (Z-value 2.734, *p*<0.05) and removed it from subsequent analyses (Supplementary Figure 3).

While samples with less than 10 samples were useful for confirming that the expected dominant taxon was present (Rickettsiaceae), we recognized that pools with less than 10 samples might not accurately reflect the diversity within populations. We also wanted to compare the effect of sex on microbial diversity and could not do so with pools containing DNA from both males and females. Therefore, we focused all subsequent analyses on pools with at least 10 samples and on pools that were either all male or all female.

The dominant eubacterial families identified included Acetobacteriaceae, Anaplasmataceae, Aeromonadaceae, Bacillaceae, Burkholderiaceae, Coxiellaceae, Enterobacteriaceae, Flavobacteriaceae, Rickettsiaceae, and Spirochaetaceae (Figure 2A). By far the most abundant were members of the Rickettsiaceae *(Rickettsia)* and Coxiellaceae *(Rickettsiella/Diplorickettsia).* Because the Rickettsiaceae were so abundant, it was difficult to see the other less-abundant taxa, so for visualization purposes, we removed Rickettsiaceae in Figure 2B. However, they were not removed from the downstream analyses. Reads matching Anaplasmataceae were mostly from genera *Wolbachia,* but also included *Anaplasma. Elizabethkingia* accounted for most of the Flavobaceriaceae, although some OTUs matched the genus *Flavobacterium.*

**Figure 2.**
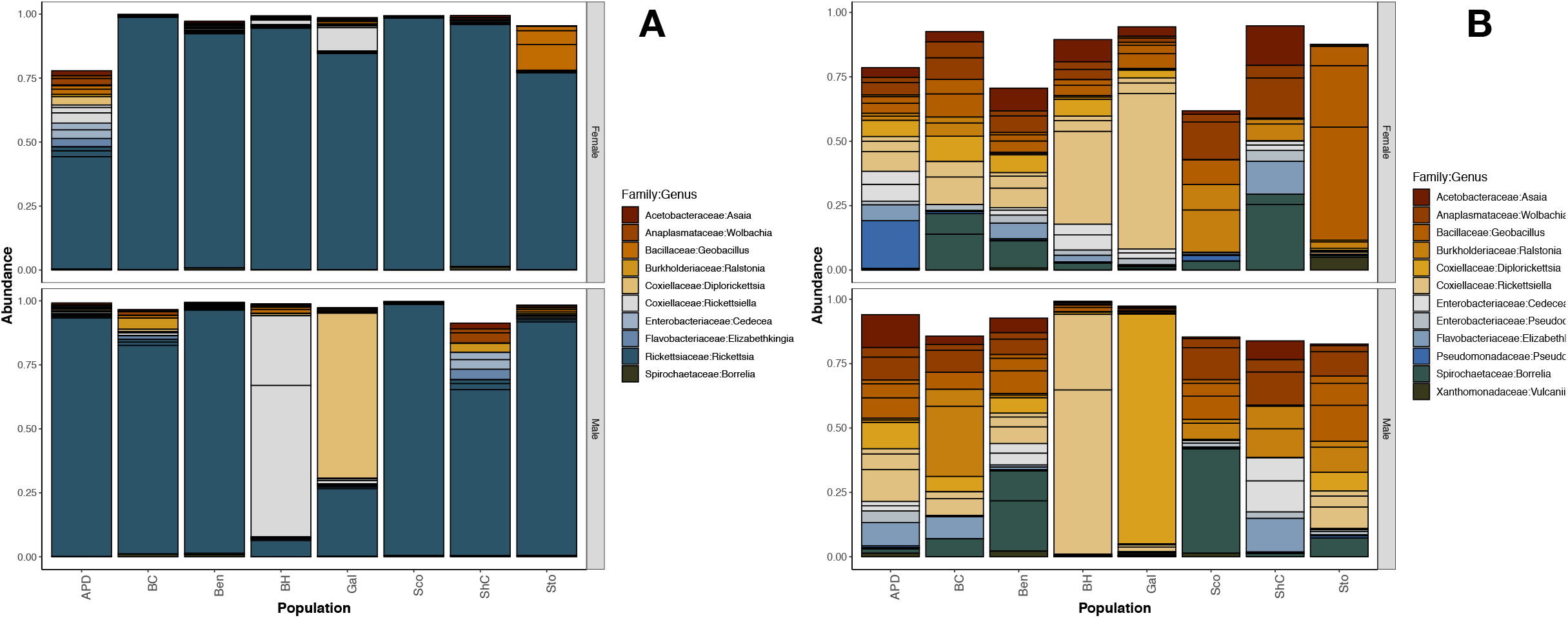
Relative read abundance of 16S rRNA (hypervariable region v4) sequenced from pools from PA. Plots show the most abundant bacterial families sorted by population and sex. Pools shown have more than 10 individuals and contain only male or only female ticks. A. With Rickettsiaceae. B. Without Rickettsiaceae. The taxonomic assignment was done using the Silva (v128) reference database.

Diversity indices suggest that female pools clustered tightly, while male pool diversity was more diffused (variable), but not significantly (Figure 3). In the principal coordinates analysis, much of the variability could be accounted for across two axes (51.8% x 18.8%) (Figure 4). We did not detect a significant effect of location but did find a significant difference between sexes (permanova: 999 permutations, *p*=0.019).

**Figure 3.**
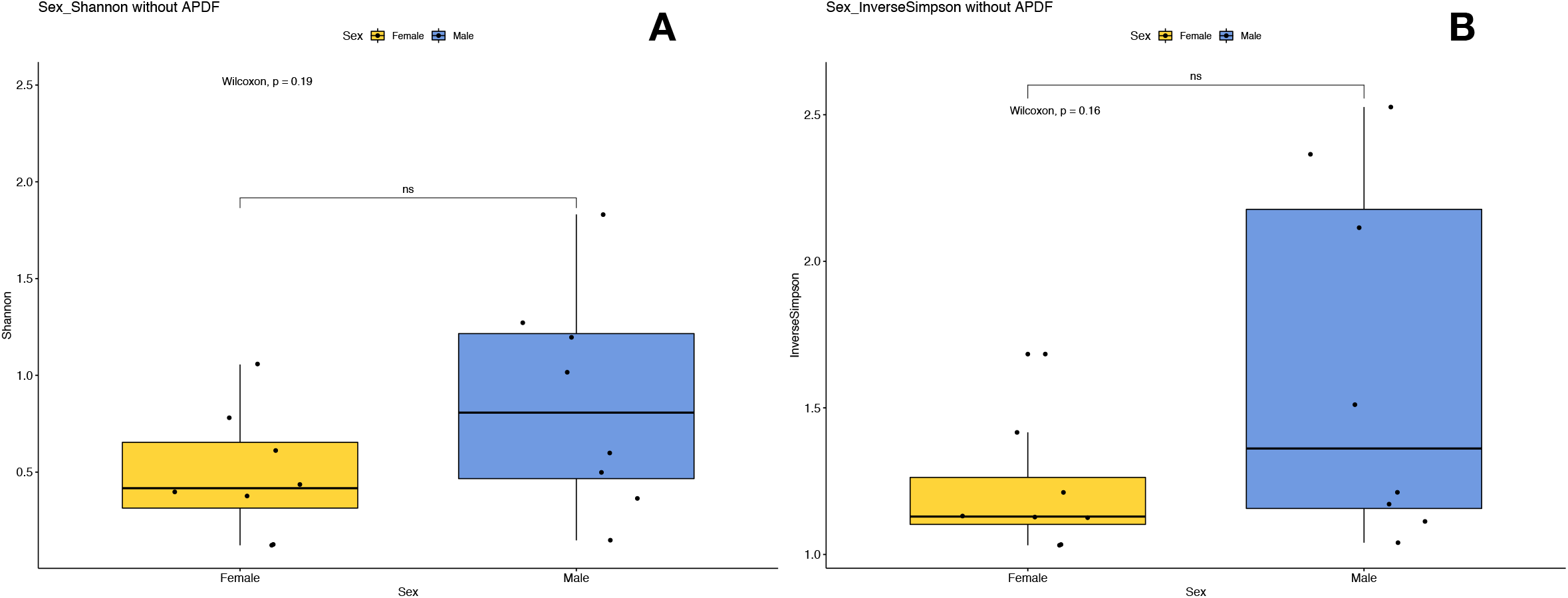
Plots of Alpha diversity indices of females versus males PA populations with at least 10 individuals. Plot A represents the Shannon index and plot B represents Inverse Simpson. The spread of indices between males pools is greater than the spread between female pools, but not significantly.

**Figure 4.**
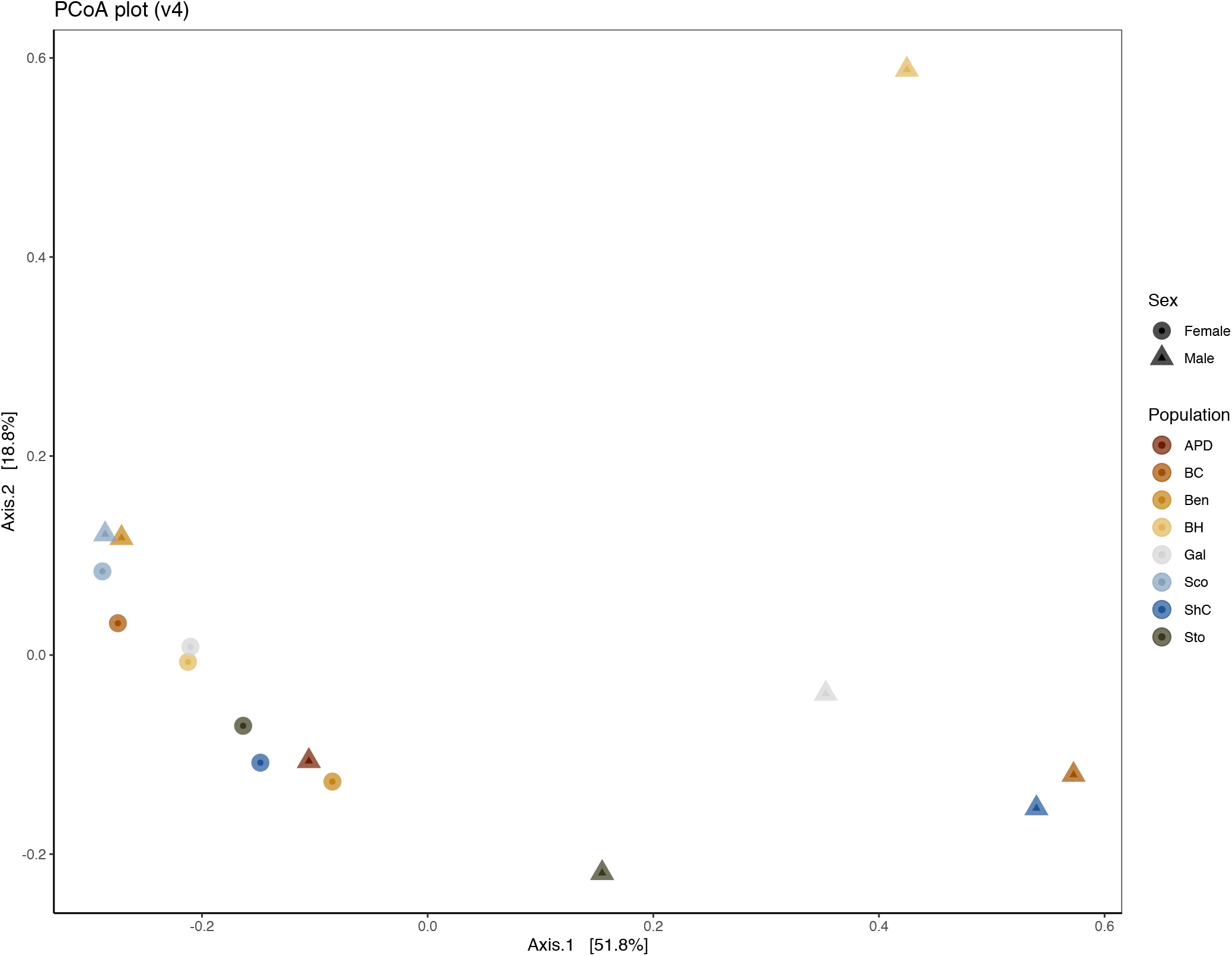
Principal Coordinate Analysis (PCoA) plot of 16S rRNA data from pooled samples of sample-based ecological distance. Colors represent populations, while shapes indicate sex. Female pools cluster together while male pools were more diverse.

### Confirmation by gene-specific amplification and sequencing

In all individuals and pools, we found the most abundant taxon (with two exceptions) was an alpha-proteobacterium in the family Rickettsiaceae, genus *Rickettsia.* Fragments from each of these pools were amplified with taxon-specific primers, sequenced, and aligned. We determined that the *Rickettsia* outer membrane rompA fragments blast aligned to the symbiotic species *Rickettsia buchneri* at 99.56-99.81% identity across 100% of the 1595 bp fragment. Phylogenetic comparison further supported that these fragments were clustered with *R. buchneri* (Supplementary Figure 4).

In two populations (MB007 and MB010, corresponding to pools of males from Big Hollow and Galbraith, respectively), the dominant reads matched the genera *Diplorickettsia* and *Rickettsiella* (gamma-proteobacterial family Coxiellaceae). To confirm the identity, we sequenced fragments of the rpoB gene using primers to the Order *Legionellales* and found the closest match for MB007 was to *Rickettsiella melolonthae* (89.85\% of a 570 bp fragment) and the closest match for MB010 was *R. viridis* (81.50\% of 575 bp). Using a maximum likelihood analysis of similar sequences, we found that both sequences clustered with *R. viridis* and *Rickettsiella* sequences from other arthropods including ticks (Supplementary Figure 5).

A 1075 bp fragment of the *Cardinium* 16S rRNA gene was amplified, sequenced, and confirmed to match a *Cardinium* found in an *I. scapularis* cell line (Accession# AB001518.1 at 99.81%. Using a maximum likelihood analysis of similar sequences, we found the sequences clustered with *Cardinium* from other arthropods including the aforementioned cell line (Supplementary Figure 6).

### Confirmation of Borrelia burgdorferi identification and frequency in ticks tested

The 16S rRNA sequencing results of the pools indicated that all pools were infected with an OTU matching the genus *Borrelia.* We confirmed the identity of *B. burgdorferi* sensu stricto by amplifying and sequencing fragments with *B. burgdorferi-specific* primers (OspC and FlaB). Sequences for OspC (333 bp) matched *B. burgdorferi* WI91-23 plasmid WI91-23_cp26 (Accession# CP001446) at 100% identity. The FlaB sequences (466 bp) matched *Borrelia burgdorferi* strain B31_NRZ (accession# CP019767.1) at 100% identity. The frequency of infected individuals per population was between 20 and 75%)(Figure 5). There was no significant difference in *B. burgdorferi* abundance between male and female adult ticks: OspC (Mann-Whitney U=31.5, z-score =0.751, *p*-value = 0.453) or FlaB (Mann-Whitney U=24.5, z-score = 1.369, *p*-value = 0.171).

**Figure 5.**
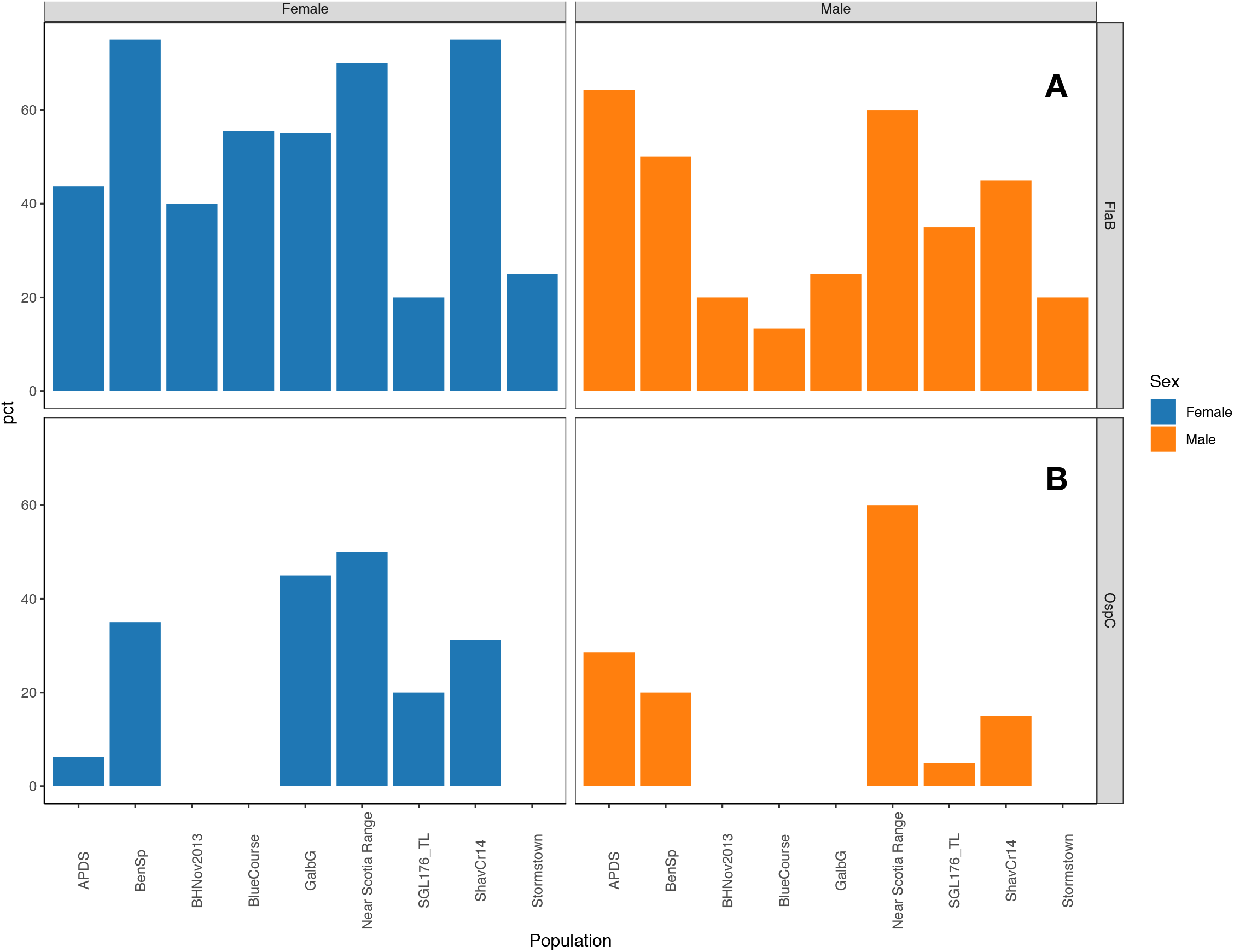
*Borrelia burgdorferi* infection frequency of individual ticks. Top chart (A) represents *B. burgdorferi* infection frequency of individuals within each pool of male or female ticks by FlaB. The bottom chart (B) represents *B. burgdorferi* infection frequency of individuals within each pool of male or female ticks by OspC.

### Relative abundance of key taxa by sex and population

We compared titers of *Rickettsia* and *Rickettsiella* by qPCR of pooled DNA sent for sequencing, additional pools used for validation, and the individuals from two pools of males with high *Rickettsiella* read counts (15 males from BH [MB007)] and 20 males from Galbraith [MB010)], and from 20 individual females from a population with low titers of *Rickettsiella* (Shaver’s Creek [MB013]). No significant titer differences for *Rickettsia* nor *Rickettsiella* were detected between sexes in pooled DNA (*Rickettsia* two-tailed Mann Whitney U= 43.5, z-score =1.85, *p*-value=0.064; *Rickettsiella* Mann Whitney U= 45, z-score = −0.985, *p*-value=0.327) (Figure 6).

**Figure 6.**
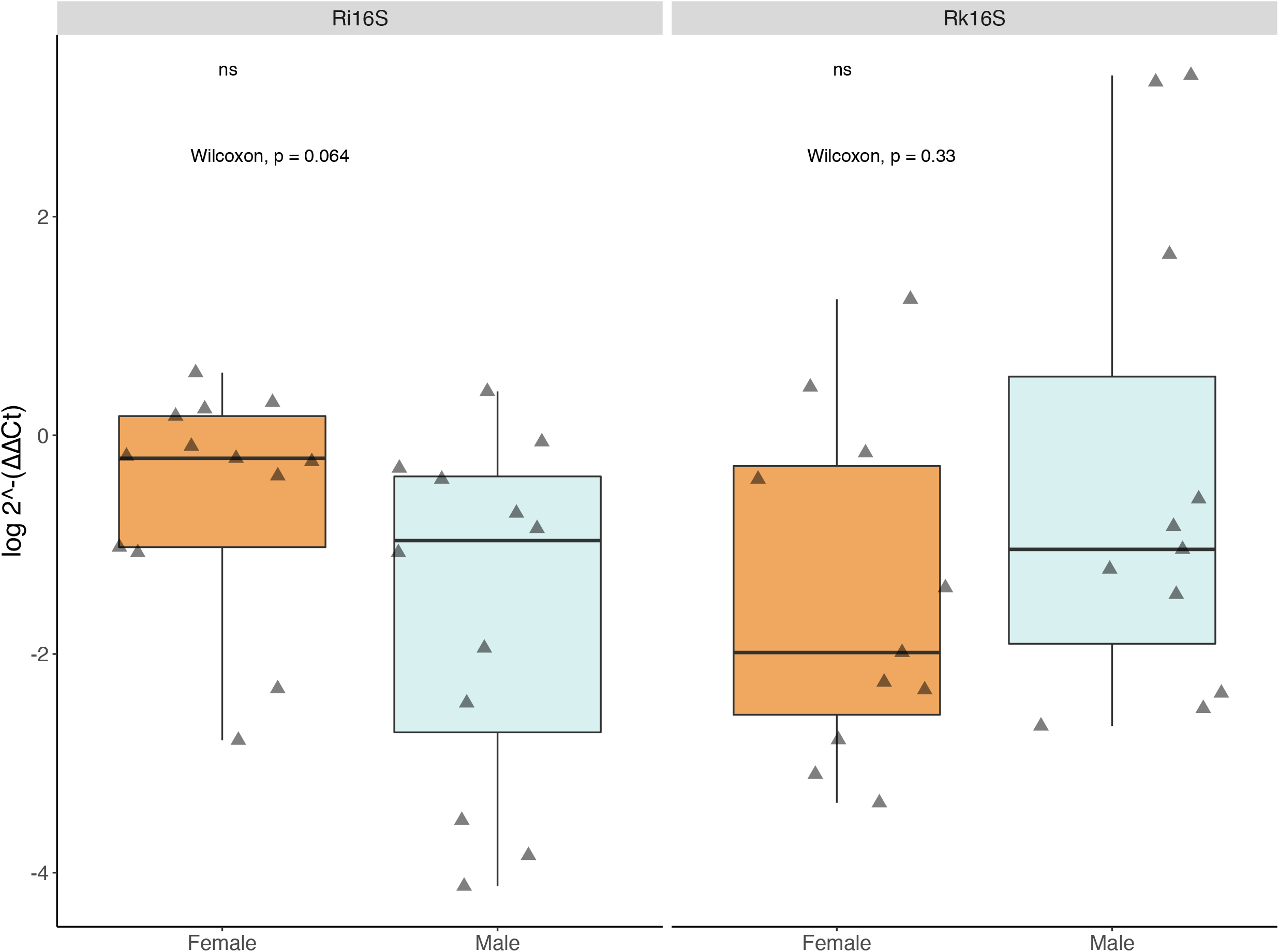
Relative titers of bacterial 16S rRNA gene from *Rickettsia* and *Rickettsiella* between pools of male and female ticks (2^-(ΔΔCt)). Data were normalized to a housekeeping gene *Ixodes scapularis* actin. Diamonds represent male or female pools tested (pools submitted for sequencing plus pools from “JulianF2015”, “BenSpF2017”, “BenSpF2018”, “HowardF” and “LedererF2019”). There For *Rickettsiella,* the pools for “JulianF2015” or “HowardF” did not amplify and were not included in the analysis. There was no significant difference between males and females in titers of *Rickettsia* nor *Rickettsiella.* Statistical significance determined by the Mann-Whitney-U calculator. (2020, January 07). Retrieved from https://www.socscistatistics.com/tests/mannwhitney/default2.aspx. Significance labels were produced in R.

When we assayed individuals from select pools of males or females, we found that *Rickettsia* titers were significantly higher in individual females versus males. (Supplementary Figure 8)(Mann-Whitney U=90.5, z-score = −4.53156, *p*-value is < .00001), while *Rickettsiella* titers in individual females were significantly lower than males (Mann-Whitney U=14.5, z-score =5.86128, *p*-value is < .00001). Within male pools, a few individual males had extremely high titers (several folds higher than others) that may account for the high read abundance of *Rickettsiella* observed in pools submitted for Illumina sequencing (Supplementary Figure 8).

We identified OTUs matching the genera Candidatus *Cardinium* and *Anaplasma phagocytophilum* at low frequencies. *Cardinium* is a known symbiotic bacteria of other arthropods in the Bacteroidetes so although the read counts for Candidatus *Cardinium* were below the threshold (not within the top 20 most abundant taxa), we chose to confirm its identity using genus-specific amplification and Sanger sequencing. In all pools assayed, only a single population matched *Cardinium,* When we screened each individual within that population, a single individual in the female pool of 20 tested positive.

We confirmed the presence and relative titers of *Anaplasma phagocytophilum* in our pools. Additionally, we tested 10 pools sorted by sex from 4 additional locations: Julian, PA (2015); SGL333 (2017 and 2018), Howard, PA (2019), and Lederer Park (2019). The ApMSP2 levels were significantly higher in male versus female tick pools (Figure 7).

**Figure 7.**
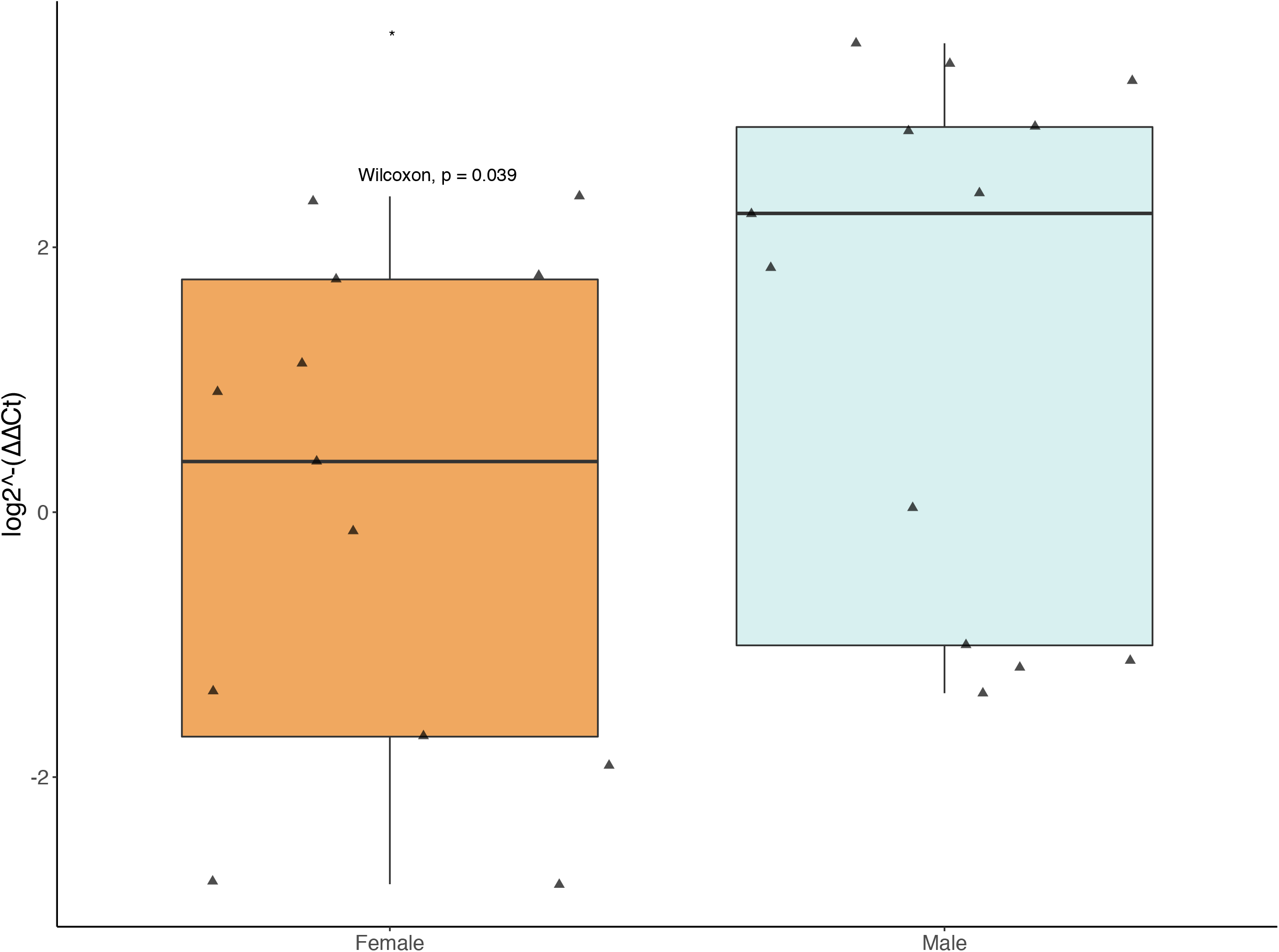
Relative titers of *Anaplasma phagocytophilum* MSP2 gene from pools of male and female ticks. Data were normalized to a housekeeping gene *Ixodes scapularis* actin. Diamonds represent male or female pools tested (pools submitted for sequencing plus pools from “JulianF2015”, “BenSpF2017”, “BenSpF2018”, “HowardF” and “LedererF2019”). Male pools had significantly higher titers of *Anaplasma phagocytophilum (*).* Statistical significance determined by the Mann-Whitney-U calculator. (2020, January 07). Retrieved from https://www.socscistatistics.com/tests/mannwhitney/default2.aspx. Significance labels were produced in R.

Sequencing results suggested that *Wolbachia* was present in almost all pools. We did not investigate the infection frequency or titers in pools, but we recognized that *Wolbachia* could be present as a symbiont of an endoparasite of the tick. We therefore tested for presence of wasp or nematode DNA, but did not detect either.

## Discussion

Tick obligate intracellular symbionts have been closely studied as potential tools for manipulation of ticks, tick-borne pathogens, or both. The first record of symbiont-pathogen antagonism in ticks was observed in populations of the wood tick *Dermacentor andersoni* from east and west sides of the Bitterroot Valley in Montana [53]. While a female could be infected dually by two closely related *Rickettsia* species, the pathogenic *R. rickettsii* was not found in ovarial tissues if the tick was also infected with the east side agent (later isolated and identified as *R. peacockii).* Researchers proposed the hypothesis of “interference”, in which a virulent pathogen was unable or less likely to invade ovarial tissues due to the presence of a less virulent or symbiotic species such as *R. peacockii* [53]. Subsequent isolation and characterization of *R. peacockii* revealed that it was genetically almost identical to the highly virulent *R. rickettsii,* but contained several insertion sequences throughout its genome [54,55]. Interference was also observed between *R. montanensis* (formerly known as *R. montana)* and *R. rhipicephalus,* and between different strains of *Anaplasma marginale* [56,57].

In this study we used 16S rRNA sequencing to determine the microbial community structure of *I. scapularis* in several populations within Centre County, PA, plus two pools from Mississippi. The most abundant reads matched the Rickettsiaceae (genus *Rickettsia),* We used rompA gene amplification and sequence comparison to confirm that this *Rickettsia* was the *I. scapularis* symbiont *R. buchneri.* Both pathogenic and nonpathogenic *Rickettsia* occur in association with multiple tick species. While microbial ecology surveys over 20 years have detected *Rickettsia* in *I. scapularis* populations throughout the eastern United States, it was not until 2015 that it was determined that all these isolates were the same species and formally named *Rickettsia buchneri* sp. novo [58]. It was therefore unsurprising to find this taxon in high abundance. *R. buchneri* is a nonpathogenic member of the spotted fever group rickettsia and is known to be transmitted trans-generationally [58]. *R. buchneri* contains several genes that may supplement an incomplete heme biosynthesis pathway in *I. scapularis [59].* Whether or not *R. buchneri* is essential for tick survival, plays a substantial role in defining in the microbial community, or interacts in some way with invading pathogens remains to be seen.

*Rickettsia* abundance was consistently high in most females, but was more variable in pools of males and tended to occur at lower abundance when other taxa were more abundant. Specifically, we detected OTUs matching Coxiellaceae that occurred in inverse abundance to *R. buchneri* in some pools of male ticks. At the genus-level, most pools contained reads matching the genus *Rickettsiella,* but one male pool (GalM) contained reads bioinformatically identified as *Diplorickettsia.* However, we did not detect *Diplorickettsia* when we sequenced amplicons using Legionellales-specific primers. The taxonomic status of *Diplorickettsia/Rickettsiella* seems to be somewhat dynamic. *Diplorickettsia* was designated a new genus, then subsumed back under *Rickettsiella,* but according to the Genome Taxonomy Database (gtb.ecogenomic.org), *Diplorickettsia* is not only a standalone genus but is in its own Order and Class (Diplorickettsiales: Diplorickettsiaceae), and *Rickettsiella* are contained within [43,60]. For this study, we chose to use the NCBI taxonomy, which lists both genera under Legionellales: Coxiellaceae.

*Rickettsiella* has been described in *I. scapularis* from Massachusetts, and in other *Ixodes* species [61,62]. Whether or not this particular taxon interacts with *Rickettsia* at the cellular or organismal level is not known, but it is known that *Rickettsiella* is associated with malpighian tubules and ovaries in *I. woodi* [62]. In our qPCR evaluation of the individuals from these Coxiellaceae-dominated pools, we found that the abundance was largely due to contributions of 2-3 heavily infected individuals, while the majority of individuals within these pools were heavily populated with *Rickettsia.* To date, *I. scapularis*-associated *Rickettsiella* is not known to be pathogenic to humans, but other species can be pathogenic or mutualistic in their invertebrate hosts [63]. *Rickettsiella* in *I. woodi* was hypothesized to be involved in the parthenogenic nature of one lab colony [60,62]. Presumably the *Rickettsiella* in *I. scapularis* does not cause parthenogenesis since we detected this genus in both males and females from field populations. We are investigating what, if any, effect *Rickettsiella* may have on *I. scapularis* biology.

All sex-population pools contained reads corresponding to the Spirochaetaceae matching the genus *Borrelia. Borrelia miyamotoi* has been shown to occur at low frequencies in some populations of *I. scapularis* [64], but we did not investigate this and only assessed the presence of *B. burgdorferi.* We found that 20-75% of individual ticks within each sex-population pool were infected with *B. burgdorferi,* but did not detect a sex-specific pattern.

We identified two potential genera of interest identified within the Anaplasmataceae: *Wolbachia* and *Anaplasma. Wolbachia* is an alpha-proteobacterial intracellular symbiont that infects a wide range of invertebrate hosts. *Wolbachia* can have little effect on its host or can alter its reproductive biology in different ways ranging from male-killing, parthenogenesis, or cytoplasmic incompatibility [65–67]. While this was not our experimental focus, it was an interesting observation given that *Wolbachia* has been found to positively or negatively affect pathogen transmission in other vector systems [8]. *Wolbachia* has been found in other *I. scapularis* studies [23,61], but it has not been determined whether the ticks themselves are infected or if they are parasitized by something that is itself infected with *Wolbachia.* We tested for presence of wasps and nematodes and found no evidence of either in our populations. However, without a more in-depth study, we cannot conclusively state that the *Wolbachia* identified is an infection of the ticks or an infection of an endoparasite infecting the ticks. If the incidence of nematode parasitism/wasp parasitoidism was low, we may not have tested enough samples or locations to successfully detect it. It would be interesting to determine if endoparasites are present and if so, how prevalent they are in *I. scapularis* populations.

The other identified Anaplasmataceae of interest was *Anaplasma phagocytophilum (Ap),* the etiological agent of human anaplasmosis. Cases of human anaplasmosis have been increasing since it became a notifiable disease in 2008. While some increase was expected due to increased awareness and testing, the number of reported cases has risen from 4151 in 2016 to 5762 in 2017 (https://www.cdc.gov/anaplasmosis/stats/index.html). In the Illumina data from pooled DNA from individual ticks, we identified low-titer *Ap* infections in several pools. We used Ap-specific Taqman qPCR and found a statistically higher titer of *Ap*-infected male pools versus female pools, but it is unclear what the biological significance, if any, of this would be. Males are not considered epidemiologically important as vectors. However, if they were infected, they represent infected individuals that acquired infections either as larvae or nymphs. Cases of reported human anaplasmosis cases increased during the summer months, coincident with the active period of *I. scapularis* larvae and nymphs (https://www.cdc.gov/anaplasmosis/stats/index.html). Infected adult ticks, therefore, reflect the presence of the pathogen circulating in the tick populations in the previous year.

We explored the use of pooled sequencing as an initial assessment of the core microbial community composition at the population level. The advantage of using this approach is twofold. First, we can go back to the archived samples and extract additional nucleic acids. Second, we minimize the number of samples initially submitted for sequencing to identify populations of interest for further individual-level screening. One obvious problem with this approach, though, was that individuals with high infection loads might suggest a high infection load overall, so this needed to be confirmed through assays of individuals.

The pooled approach allowed us to find a rare *Cardinium* symbiont that we confirmed to be from only one of 40 individuals tested from a single population. To our knowledge, this taxon has not been found in other next-generation sequencing-based microbiome studies, presumably because it is so rare. We do not know how widely this *Cardinium* species is distributed, nor what its effect, if any, on its host. The first published account of the *I. scapularis*-associated *Cardinium* (then referred to as the *“Cytophaga-like* organism”) was from ticks collected in Nantucket MA where it was extracted from ticks and grown in tick cells [68]. It remains to be seen whether it is a transient introduction into the local tick population (e.g. from a tick was transported on a migrating bird or mammal), whether it is affected in some way by the native microbiota, or whether it has any impact on the tick host. *Cardinium* is an intracellular bacterium known to be associated with ovaries and midguts of arthropods, so it is conceivable that it may interact with other intracellular bacteria, pathogenic or otherwise [69].

We observed several limitations of relying on Illumina data alone without a biological context. First, we realize that the number of individuals tested was relatively small for some of the populations. This was a reflection of the abundance in sampled areas, but a few individuals with heavy infections of one or more taxa can skew the overall outcomes, resulting in a misleading conclusion. Second, we found that populations that were close together geographically could have very different microbial communities (either contain unique taxa or very different abundance of taxa). Had we randomly selected one or two populations for comparison with other populations in another geographic location (e.g. another state), we might conclude that these two populations represented the diversity from the state of Pennsylvania. This would have been inaccurate and missed key taxa. Lastly, choosing the right hypervariable region and correct reference database can profoundly affect the results. In our case, this resulted in a significant setback that included re-sequencing, reanalysis, and qPCR validation. Had we not also validated these data by qPCR, we would have made an erroneous interpretation of the dominant taxa and the potential implications of tick bacterial community dynamics. Thus, although next-generation sequencing allows researchers to obtain a deeper depth of coverage, it does not account for unknown biases (v6 was not known to be biased in 2013, when the sequencing was initiated).

We still lack an understanding of how (or if) a succeeding dominant taxon can cause the reduction in a native rickettsial symbiont (e.g. through direct competition for resources or receptors or indirectly inducing host immunity or production of toxins that inhibit rickettsial intracellular growth). Why are only some individuals in certain populations or sex more heavily infected with *Rickettsiella* than *R. buchneri?* Is there direct interaction between microbes that influences the within-tick microbial community composition? Are the microbial communities pre-determined by the mother or is there any transmission of bacteria from the males to offspring? Are there bacteria transmitted from the mother to the offspring from other means (e.g. egg smearing)? Could any of these taxa be targeted for control or alternation of pathogen transmission? Whether these taxa likewise interfere with or enhance transmission of pathogens/parasites (e.g. *Anaplasma, Borrelia, Babesia)* also remains to be seen.

## Conclusion

It is important to remember that, while bacterial 16S sequencing is a powerful tool for exploring taxa present in a given study, it is merely a basis for generating hypotheses and should not be relied upon to extrapolate conclusions without subsequent validation of infection frequencies. We targeted specific taxa from pools or individual tick DNA the individual tick data to determine titers or infection frequencies. With the caveat that our data represent a small sample size, our data suggest that the microbial population dynamics can be highly variable among individuals in the same population, and between populations that are at most 26 miles apart. Thus, given that diversity at the local scale is so variable, patterns between and across large geographic areas should be considered suspect without sufficient sampling of each chosen population and across the collection range.

## Author contributions

JMS collected and processed field samples, conducted molecular experiments, designed primers, optimized and ran qPCR assays, conducted analyses, wrote the manuscript, and handled revisions. GES contributed to processing tick samples and determining *Borrelia burgdorferi* infection frequency in each population. EAW contributed to field collections, processing ticks, and running PCR assays. All authors read and approved the final manuscript.

## Acknowledgements

The authors would like to thank Dr. Heather Hines, Dr. Andrew Deans, Dr. Ralph Mumma, Dr. Diana Cox-Foster, Tracey Bessemer, and David Love for allowing us to collect ticks on their property. The authors would also like to thank Dr. Shelby Fleischer, Rebecca Johnson, Dr. Duverney Chaverra-Rodriguez, Dr. Hitoshi Tsujimoto, Dr. Levent Aydin, Chaz Bunce, and Justine Alexander, who volunteered to collect ticks. The authors would like to thank Dr. Bing Ma, Brittany Dodson, and Dr. Jason Rasgon for insightful discussion and anonymous reviewers for critical manuscript review.

