## Supplementary figures and images for "Microbial ecology of *Ixodes scapularis* from Central Pennsylvania, USA"

### FigureS1.pdf

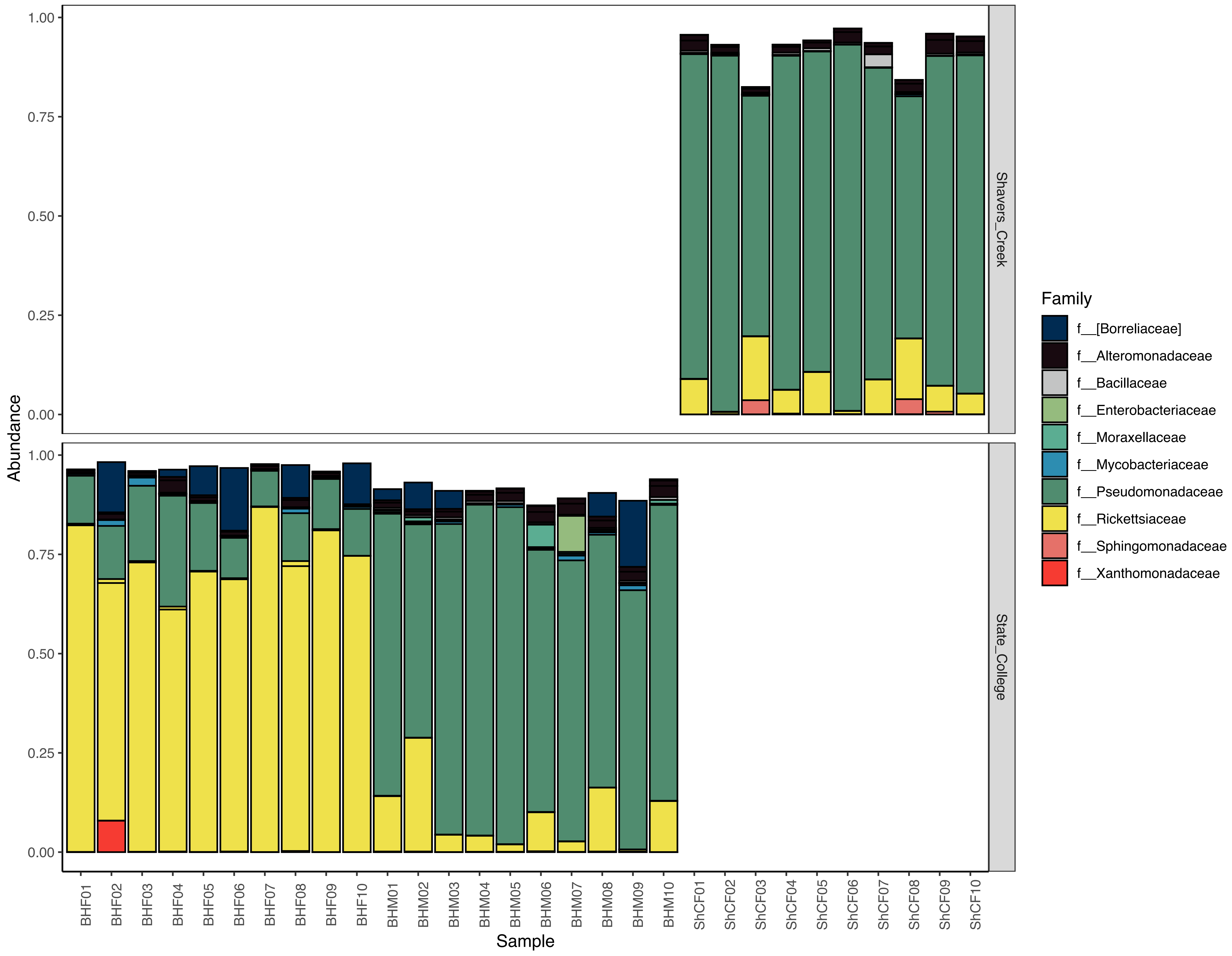

### FigureS3.pdf

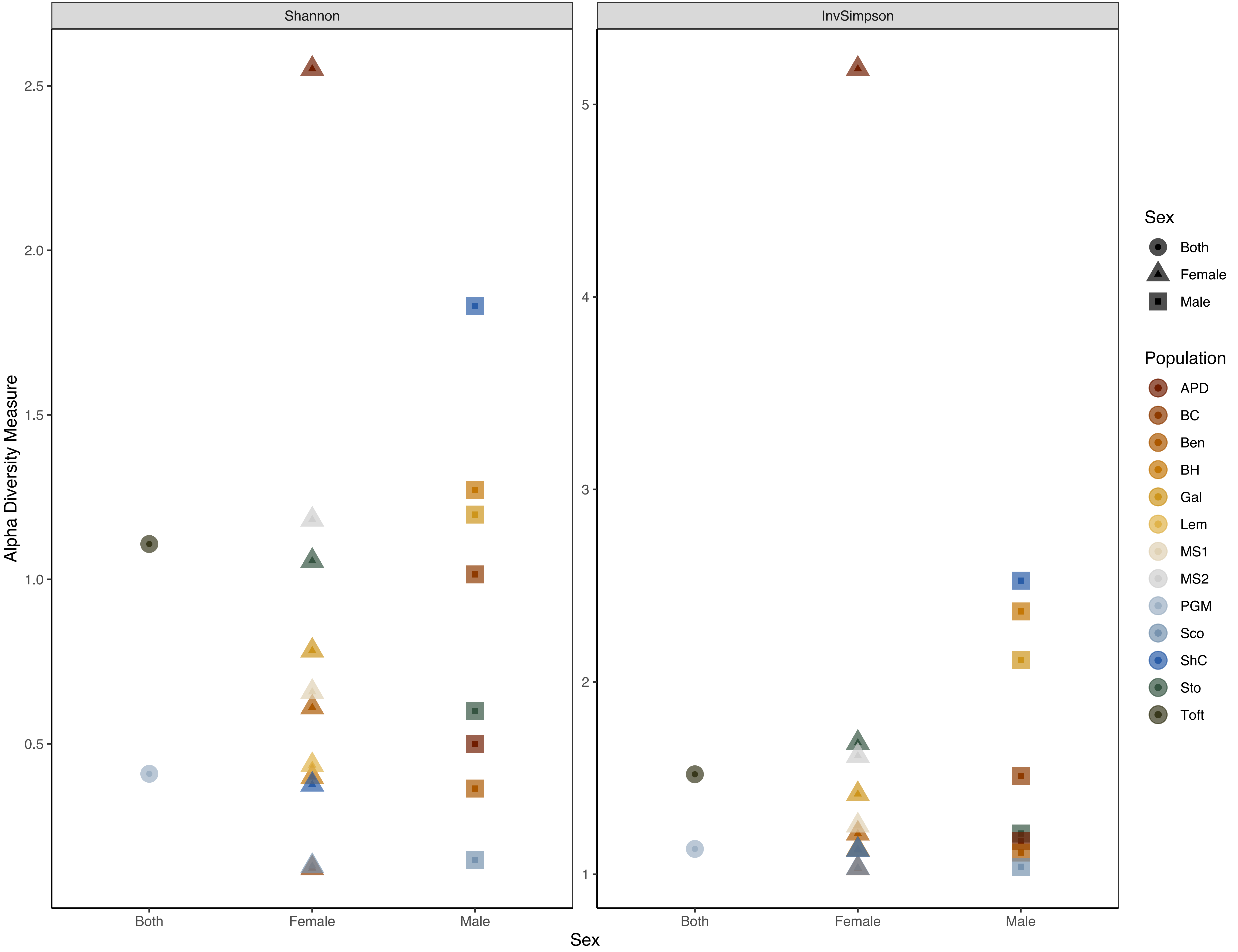

### FigureS4.pdf

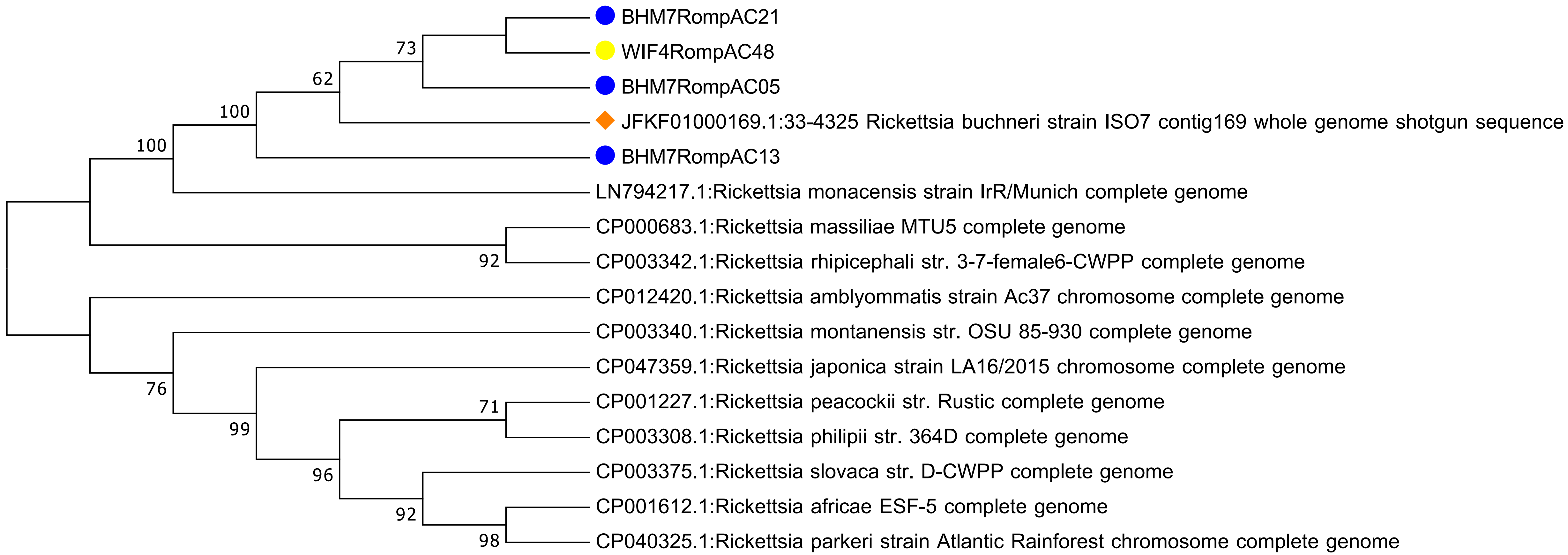

### FigureS5.pdf

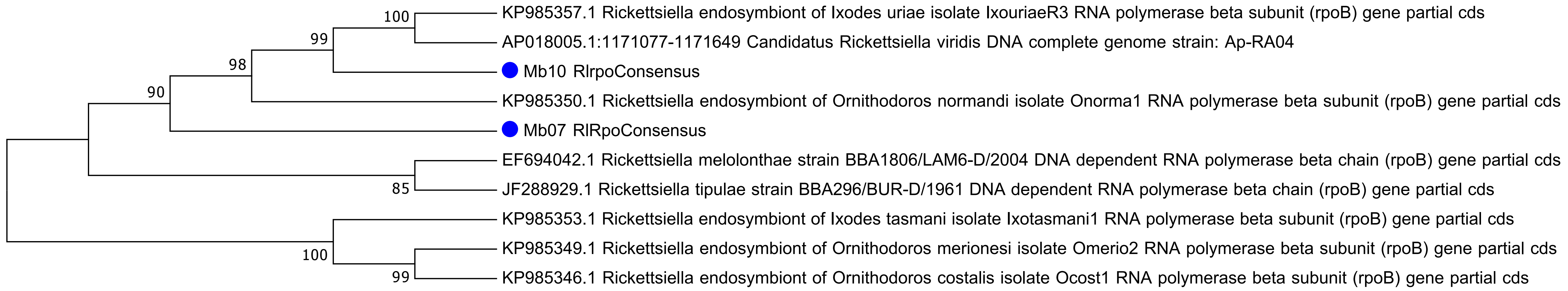

### FigureS6.pdf

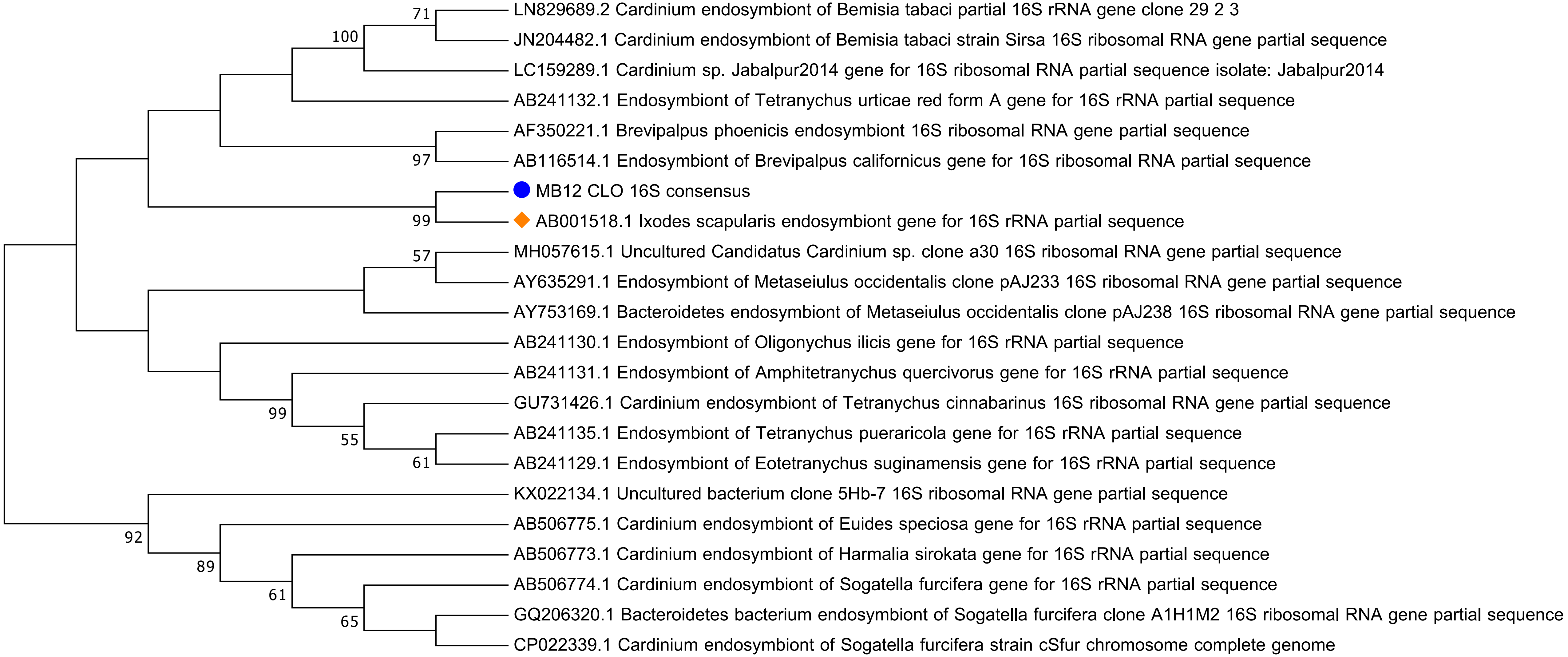

### FigureS7.pdf

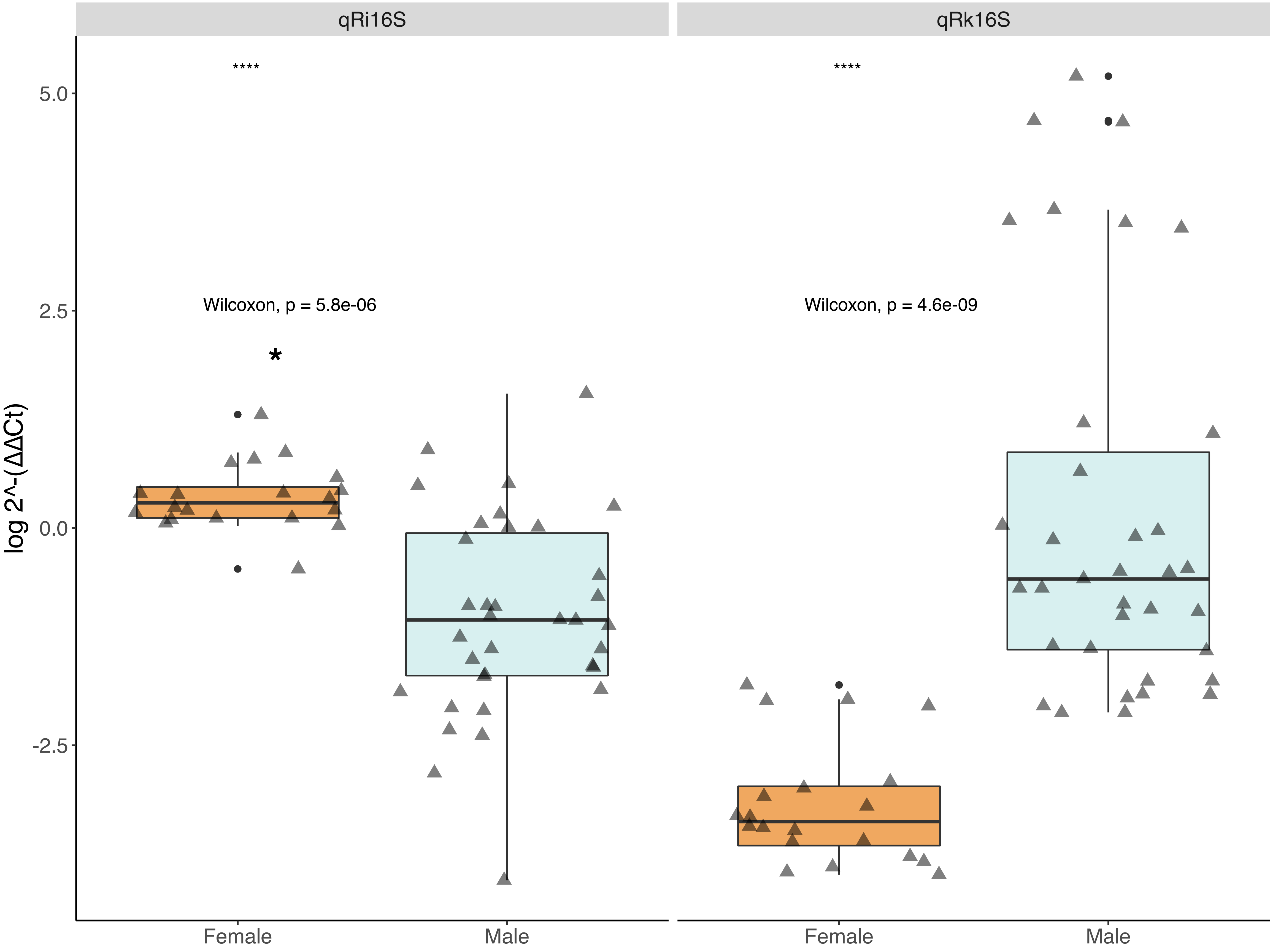
