## Supplementary material for "Microbial ecology of *Ixodes scapularis* from Central Pennsylvania, USA": FigureS2: FigureS2.pdf

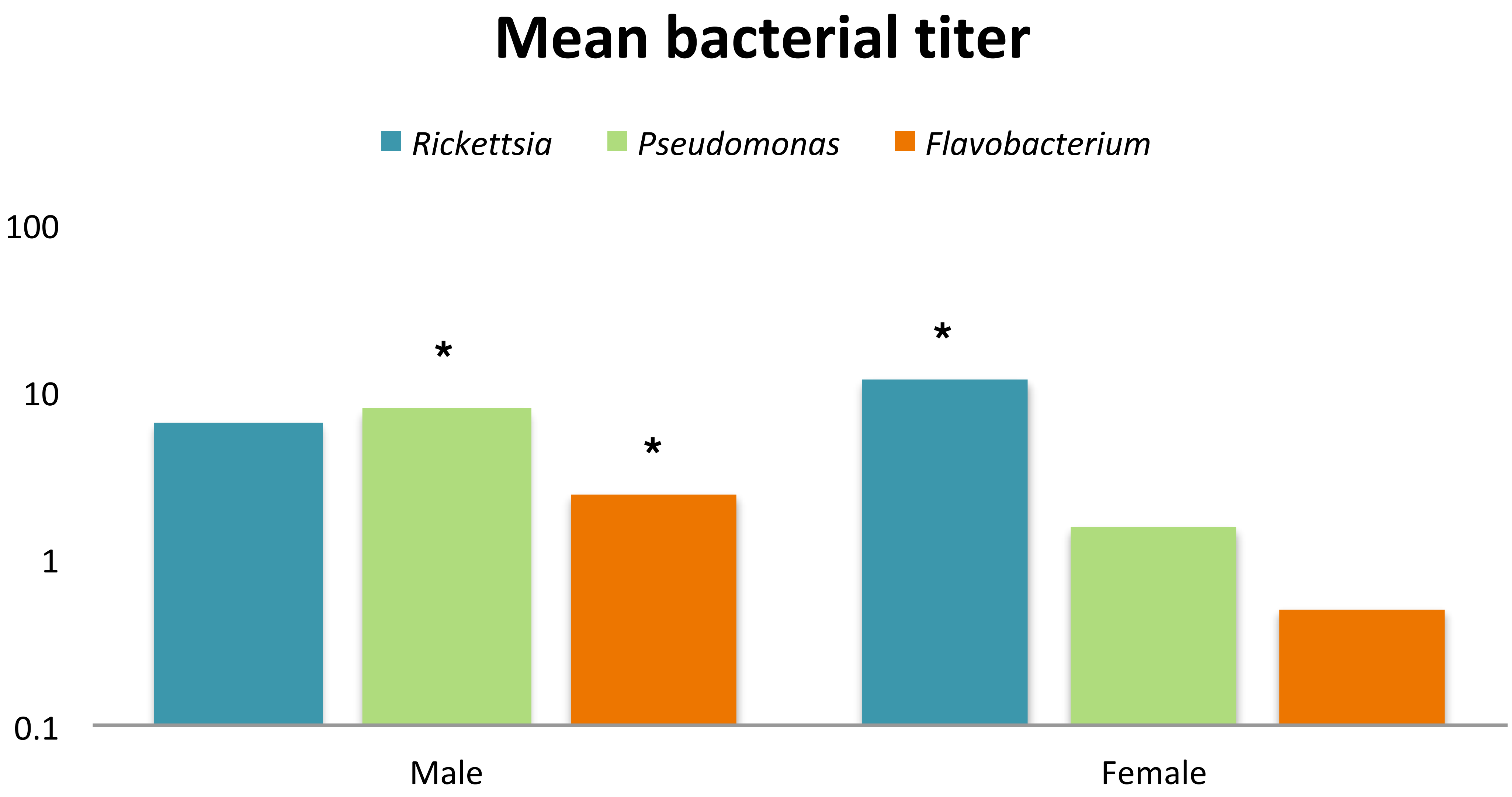


*Rickettsia*: Mann Whitney U= 87.5,  z-score = 4.66816, p-value < .00001.

*Pseudomonas*: Mann Whitney U= 116.5,  z-score = -4.15146, p-value < .00001.

*Flavobacterium*: Mann Whitney U= 113.5,  z-score = -4.20491, p-value < .00001.
